# Concentration-dependent suppression of macrolide resistance in *Mycobacterium avium* by combination therapy: an improved *in vitro* time-kill assay and PK/PD modelling study

**DOI:** 10.64898/2025.12.21.695645

**Authors:** Fumiya Watanabe, Yuta Morishige, Akio Aono, Yuji Morita, Kozo Morimoto, Satoshi Mitarai, Kazuhiko Hanada

**Affiliations:** Department of Pharmacometrics and Pharmacokinetics, Meiji Pharmaceutical University, Tokyo, Japan; Department of Pharmacy, Fukujuji Hospital, Japan Anti-Tuberculosis Association, Tokyo, Japan; Research Institute of Tuberculosis, Japan Anti-tuberculosis Association, Tokyo, Japan; Department of Infection Control Science, Meiji Pharmaceutical University, Tokyo, Japan; Respiratory Disease Center, Fukujuji Hospital, Japan Anti-Tuberculosis Association, Tokyo, Japan

**Author notes:** **Corresponding Author**: Fumiya Watanabe, Department of Pharmacometrics and Pharmacokinetics, Meiji Pharmaceutical University, 2-522-1, Noshio, Kiyose City, Tokyo 204-8588, Japan.

**Keywords:** Non-tuberculosis mycobacteria, Mycobacterium avium, PK/PD, Pharmacokinetics, Pharmacodynamics, Dose optimization

## Abstract

**Objectives:** Macrolides are key drugs in the treatment for *Mycobacterium avium* pulmonary disease, and combination chemotherapy is essential to preventing macrolide resistance. However, the concentrations of concomitant agents required to suppress resistance emergence remain undefined. This study aimed to quantify the concentration-dependent effects of companion drugs on macrolide resistance using an improved time-kill assay and pharmacokinetic/pharmacodynamic (PK/PD) modelling approach.

**Methods:** Time-kill assays were performed on *M. avium* ATCC 700898 using azithromycin alone or in combination with ethambutol, rifampicin, amikacin, or clofazimine. Total and resistant bacterial populations were quantified over 28 days. A pharmacodynamic (PD) model describing the dynamics of susceptible and resistant subpopulations was linked to a pharmacokinetic (PK) model incorporating alveolar macrophage concentrations to simulate resistance and bactericidal rates under various dosing regimens.

**Results:** Macrolide resistance most frequently emerged at 2–4× the minimum inhibitory concentration (MIC) of azithromycin, whereas resistance was less frequent at 8–16× MIC or sub-MIC levels. The addition of companion drugs suppressed resistance emergence even at sub-MIC levels. PK/PD simulations demonstrated that standard-dose ethambutol effectively prevented macrolide resistance, whereas rifampicin provided minimal protection. More frequent dosing schedules enhanced resistance suppression compared with once-daily regimens.

**Conclusion:** This study defines the concentration-dependent contributions of concomitant agents to macrolide resistance suppression in *M. avium* and establishes a quantitative PK/PD framework for evaluating resistance emergence. These findings provide mechanistic insights to support the optimization of combination dosing strategies for *M. avium* pulmonary disease.

## INTRODUCTION

The global incidence and prevalence of nontuberculous mycobacterial (NTM) pulmonary disease are increasing, and *Mycobacterium avium*-*intracellulare* complex (MAC) accounts for the majority of these NTM infections.^1^ Macrolides are key drugs for the treatment of MAC pulmonary disease, and their *in vitro* susceptibility correlates closely with the clinical outcomes.^1,2^ Therefore, current treatment guidelines recommend the use of combination chemotherapy with agents such as ethambutol (EMB), rifampicin (RIF), and amikacin (AMK) to prevent the emergence of macrolide resistance.^1^ MAC pulmonary disease predominantly affects older adults, and since the treatment duration often exceeds 12 months,^1^ adverse events leading to treatment discontinuation occur in 9.1–38% of the cases.^3–5^ In the treatment of MAC pulmonary disease, there are limited alternative antibiotic options, and no established criteria exist for dose reduction; therefore, the suspected causative agent is often discontinued when adverse events occur. Consequently, the global adherence to guideline-based therapies ranges from 8–42%.^6,7^ Deviation from the recommended regimen is a known risk factor for treatment failure and development of macrolide resistance.^8,9^ Therefore, maintaining treatment continuity through appropriate management of adverse events is crucial. However, the relationship between the dosing regimens of concomitant antimicrobials and the emergence of resistance remains unclear.

Antibiotic concentration in the epithelial lining fluid (ELF) is an important determinant of appropriate dose optimization.^10^ Specifically, intracellular antibiotic concentrations within alveolar macrophages (AM), where mycobacteria reside and survive, are critical factors in dose optimization for the treatment of MAC pulmonary disease.^11^ The concentration-time profiles of antimicrobial agents in ELF and AM have been well-characterized,^12^ allowing the estimation of mycobacterial drug exposure at the site of infection. However, to our knowledge, no studies have determined the drug concentrations required to suppress the emergence of macrolide resistance. Therefore, this study aimed to identify the concentrations of concomitant antimicrobials that suppress macrolide resistance using an improved *in vitro* time-kill assay (TKA) developed for combination therapy, and to simulate the relationship between antimicrobial dosing regimens and emergence of resistance through pharmacokinetic/pharmacodynamic (PK/PD) modelling analysis. Our findings will help optimize combination dosing strategies for *M. avium* pulmonary disease.

## METHODS

### Antibiotics, Medium, and Bacteria

Clarithromycin (CLR) and clofazimine (CFZ) were purchased from Tokyo Chemical Industry Co. Ltd. (Tokyo, Japan). Azithromycin (AZM) and EMB were purchased from LKT Laboratories, Inc. (MN, USA). RIF and AMK were purchased from FUJIFILM Wako Pure Chemical Co. (Osaka, Japan). Middlebrook 7H9 broth (MB7H9b), cation-adjusted Mueller-Hinton broth (CAMHb), Middlebrook 7H10 (MB7H10) agar, and oleic albumin-dextrose-catalase (OADC) growth supplements were obtained from Becton Dickinson and Co. (NJ, USA). *M. avium* ATCC 700898 (American Type Culture Collection, Manassas, VA, USA) was used for all the TKAs. The strain was stored at -80°C and was freshly cultured before each assay to ensure the use of bacteria in the logarithmic growth phase.

### Drug Susceptibility Testing

For each antibiotic, the minimum inhibitory concentration (MIC) against *M. avium* was determined according to the Clinical and Laboratory Standards Institute (CLSI) M24 3^rd^ ed recommendations,^2^ except that for CFZ, wherein we determined the MIC of MB7H9b instead of CAMHb, owing to the poor solubility of CAMHb.^13^ Additionally, we confirmed that the MICs obtained using MB7H9b were consistent with those measured using CAMHb.

### TKAs against *M. avium*

TKAs for the antimicrobials were performed using previously published methods, with modifications.^13^ Previously, TKAs for *M. avium* have focused on the overall bacterial growth under combination therapy, while this study aimed to evaluate the suppression of macrolide resistance. To this end, we first established conditions under which macrolide monotherapy consistently produced stable and frequent emergence of resistance and then conducted combination TKAs with EMB, RIF, AMK, and CFZ to assess the relationship between these concentrations and the suppression of macrolide resistance. All TKAs were performed in quadruplicate at 37°C with shaking at 135 rpm.

### TKA for Macrolide Monotherapy

Bottles containing 10 mL MB7H9b with 0.05% Tween 80 and 10% OADC were inoculated with *M. avium* at an initial density of approximately 2 × 10^7^ colony-forming units (CFU)/mL. AZM was added at concentrations ranging from 0.25× to 16× MIC. On days 0, 7, 14, 21, and 28, the number of total bacteria and macrolide-resistant bacteria was quantified using the triplicate spot-plating method on MB7H10 agar plates and 32 mg/L CLR-containing MB7H10 agar plates, respectively, according to the CLSI criteria.^2^ The plates were incubated at 37°C for 7Ld, and the CFU/mL were determined. The lower limit of detection (LOD) was 100 CFU/mL. To confirm resistance mutations at the molecular level, six colonies were randomly selected from each of the following sources: the original inoculum before the experiment, colonies appearing only on MB7H10 agar after incubation, and colonies appearing on MB7H10 agar containing 32 mg/L CLR. DNA was extracted from each colony and point mutations at positions 2058 and 2059 in the *rrl* gene region, which are responsible for macrolide resistance,^2^ were analysed using Sanger sequencing.

### TKAs under Combination Exposure

Under conditions where resistance emergence was most frequently observed in the previous experiment, the AZM concentration was fixed, and EMB, RIF, AMK, or CFZ were co-incubated at concentrations ranging from 0.03× to 2× of their respective MICs. The number of total and macrolide-resistant bacteria was quantified. To evaluate the stability of the antimicrobial agents, quality control samples at 0.25× and 16× MIC for each drug were prepared in quadruplicate and incubated with aliquots collected at 0, 7, 14, 21, and 28 d after initiation. Drug concentrations were measured using a validated high-performance liquid chromatography-mass spectrometry method, as described previously.^14^

### Pharmacodynamic Modelling

The *in vitro* TKA data were analysed using PD modelling. The details are presented in Text S1. Briefly, a PD model was developed using NONMEM version 7.6.0 (ICON Development Solutions, Hanover, MD, USA). The model comprises susceptible and resistant bacterial subpopulations, each characterized by distinct growth, antibiotic-mediated killing, and resistance acquisition rate constants. Antibiotic concentrations were corrected based on first-order degradation rate constants (K _deg_) determined from the stability of each drug.

### PK/PD Model Simulation

To simulate the risk of macrolide resistance under different dosing regimens, we developed a PK model to estimate the ELF and AM concentration profiles of the antimicrobial agents and linked it to the PD model developed in this study. Since information on CFZ concentration–time profiles in pulmonary ELF and AM was not available, CFZ was not included in the PK/PD analysis. For PK model construction, population PK parameter estimates, such as clearance and volume of distribution, which describe the plasma concentration–time profiles of each antimicrobial agent, as well as the penetration ratios into ELF and AM, were obtained from previous reports,^12,14–31^ details are provided in Text S2.

Using the developed PK/PD model, macrolide resistance rates and bactericidal effects in AM after 6 months of two-drug combination therapy, such as AZM+EMB, AZM+RIF, or AZM+AMK were simulated under different dosing intervals (three times weekly, once daily, or twice daily), doses, and renal functions (creatinine clearance). Similarly, simulations were performed for the three- and four-drug combination regimens, including AZM+EMB+RIF, AZM+EMB+AMK, and AZM+EMB+RIF+AMK. The emergence of resistant bacteria above the LOD (100 CFU/mL) was defined as resistance, and the total bacterial count below the LOD was defined as eradication. For each condition, N = 500 simulations were performed. All simulations, data summarizations, and visualizations were conducted using R version 4.5.2 (R Foundation for Statistical Computing, Vienna, Austria) using the rxode2 and ggplot2 packages.^32^

## RESULTS

The measured MICs for AZM, EMB, RIF, AMK, and CFZ were 8, 8, 0.5, 8, and 8 mg/L, respectively, which were identical to those of MB7H9b. Genome analysis revealed the absence of *rrl* point mutations in the pre-experimental strain and colonies grown on MB7H10 agar. In contrast, all colonies isolated from MB7H10 agar containing 32 mg/L CLR harboured either an A2058C or A2059G mutation in the *rrl* gene region.

### TKA analysis

Culturing with AZM monotherapy resulted in the frequent and consistent emergence of macrolide resistance at concentrations of 2–4× MIC, where almost all bacteria in the medium were replaced by resistant populations (Figure S1). In contrast, at concentrations below the MIC, the resistant bacteria did not outgrow the susceptible population. This trend was independent of the initial bacterial inoculum, since similar results were observed with initial bacterial loads of 10L and 10L CFU/mL (Figure S1).

In the TKAs performed using combination therapy, AZM was fixed at 4× MIC, a condition prone to resistance development, and co-incubated with concomitant agents at various concentrations. The concentrations of concomitant antimicrobials that completely suppressed macrolide resistance were 0.25× MIC for EMB, 0.25× MIC for RIF, 0.125× MIC for AMK, and 0.125× MIC for CFZ. In addition, RIF and AMK at 1× MIC and CFZ at 0.25× MIC eliminated all bacteria, including susceptible populations. In contrast, EMB showed no bactericidal effects against the susceptible bacteria (Figure S2). This trend was also observed at an initial bacterial load of 10 CFU/mL (Figure S3).

### PD modelling

The stability of each drug in the culture medium is shown in Figure S4. The K _deg_ values for AZM, EMB, RIF, and AMK were 0.0074, 0.015, 0.047, and 0.0041 per day, respectively. A two-compartment subpopulation model was fitted to the TKA data. The goodness-of-fit plots are shown in Figure S5, and the parameter estimates obtained from the PD model are summarized in Table 1. The observed and PD model-predicted bacterial counts are shown in Figure 1. The PD model developed in this study adequately captures the observed data.

**Table 1.**
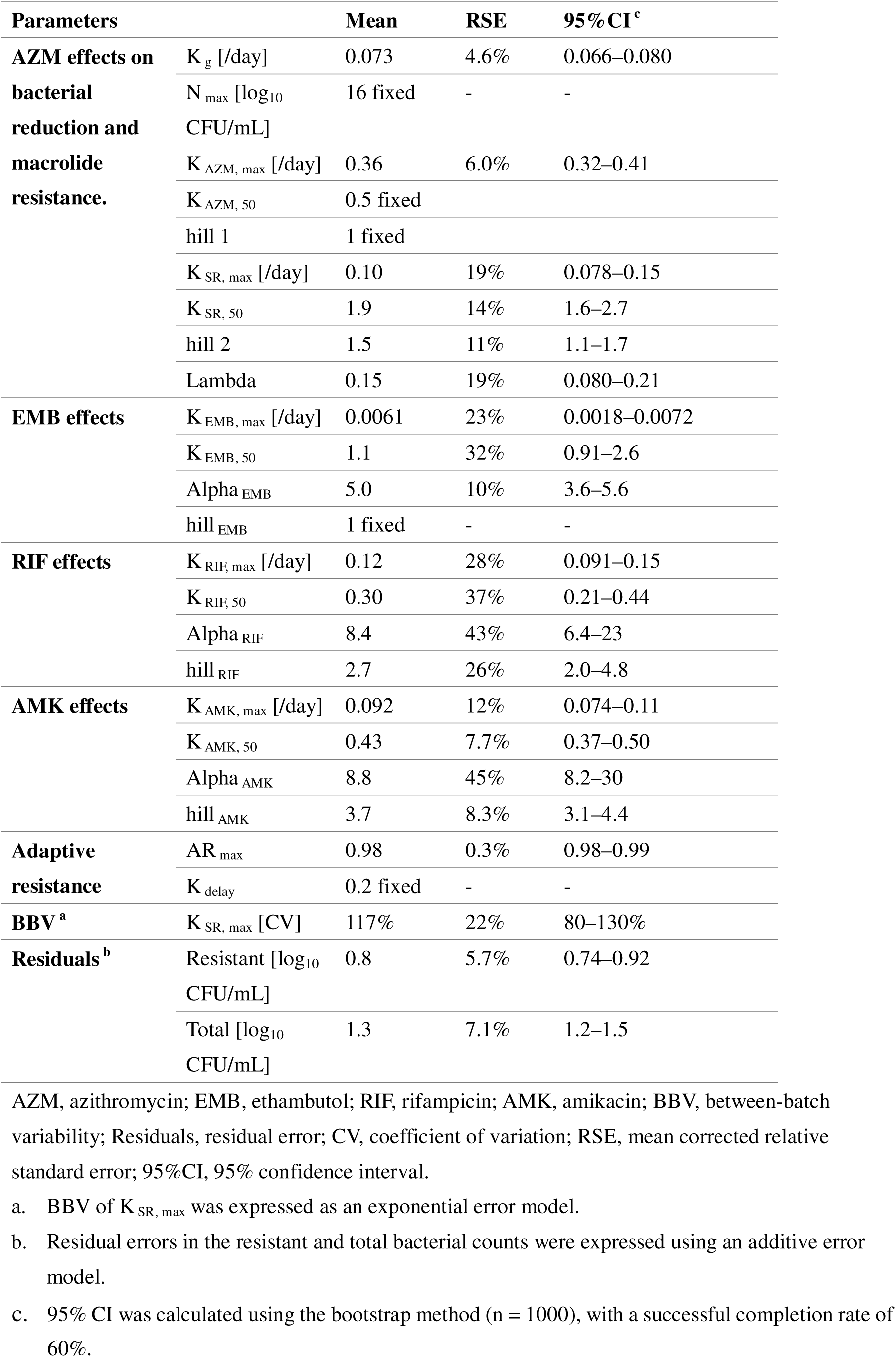
Pharmacodynamic (PD) model parameter estimates.

| Parameters |  | Mean | RSE | 95%CI<sup>c</sup> |
| --- | --- | --- | --- | --- |
| **AZM effects on bacterial reduction and macrolide resistance.** | K<sub>g</sub> [/day] | 0.073 | 4.6% | 0.066–0.080 |
|  | N<sub>max</sub> [log<sub>10</sub> CFU/mL] | 16 fixed | - | - |
|  | K<sub>AZM, max</sub> [/day] | 0.36 | 6.0% | 0.32–0.41 |
|  | K<sub>AZM, 50</sub> | 0.5 fixed |  |  |
|  | hill 1 | 1 fixed |  |  |
|  | K<sub>SR, max</sub> [/day] | 0.10 | 19% | 0.078–0.15 |
|  | K<sub>SR, 50</sub> | 1.9 | 14% | 1.6–2.7 |
|  | hill 2 | 1.5 | 11% | 1.1–1.7 |
|  | Lambda | 0.15 | 19% | 0.080–0.21 |
| **EMB effects** | K<sub>EMB, max</sub> [/day] | 0.0061 | 23% | 0.0018–0.0072 |
|  | K<sub>EMB, 50</sub> | 1.1 | 32% | 0.91–2.6 |
|  | Alpha<sub>EMB</sub> | 5.0 | 10% | 3.6–5.6 |
|  | hill<sub>EMB</sub> | 1 fixed | - | - |
| **RIF effects** | K<sub>RIF, max</sub> [/day] | 0.12 | 28% | 0.091–0.15 |
|  | K<sub>RIF, 50</sub> | 0.30 | 37% | 0.21–0.44 |
|  | Alpha<sub>RIF</sub> | 8.4 | 43% | 6.4–23 |
|  | hill<sub>RIF</sub> | 2.7 | 26% | 2.0–4.8 |
| **AMK effects** | K<sub>AMK, max</sub> [/day] | 0.092 | 12% | 0.074–0.11 |
|  | K<sub>AMK, 50</sub> | 0.43 | 7.7% | 0.37–0.50 |
|  | Alpha<sub>AMK</sub> | 8.8 | 45% | 8.2–30 |
|  | hill<sub>AMK</sub> | 3.7 | 8.3% | 3.1–4.4 |
| **Adaptive resistance** | AR<sub>max</sub> | 0.98 | 0.3% | 0.98–0.99 |
|  | K<sub>delay</sub> | 0.2 fixed | - | - |
| **BBV<sup>a</sup>** | K<sub>SR, max</sub> [CV] | 117% | 22% | 80–130% |
| **Residuals<sup>b</sup>** | Resistant [log<sub>10</sub> CFU/mL] | 0.8 | 5.7% | 0.74–0.92 |
|  | Total [log<sub>10</sub> CFU/mL] | 1.3 | 7.1% | 1.2–1.5 |

AZM, azithromycin; EMB, ethambutol; RIF, rifampicin; AMK, amikacin; BBV, between-batch variability; Residuals, residual error; CV, coefficient of variation; RSE, mean corrected relative standard error; 95%CI, 95% confidence interval.
a. BBV of K<sub>SR, max</sub> was expressed as an exponential error model.
b. Residual errors in the resistant and total bacterial counts were expressed using an additive error model.
- c. 95% CI was calculated using the bootstrap method ( $n = 1000$ ), with a successful completion rate of 60%.

**Figure 1.**
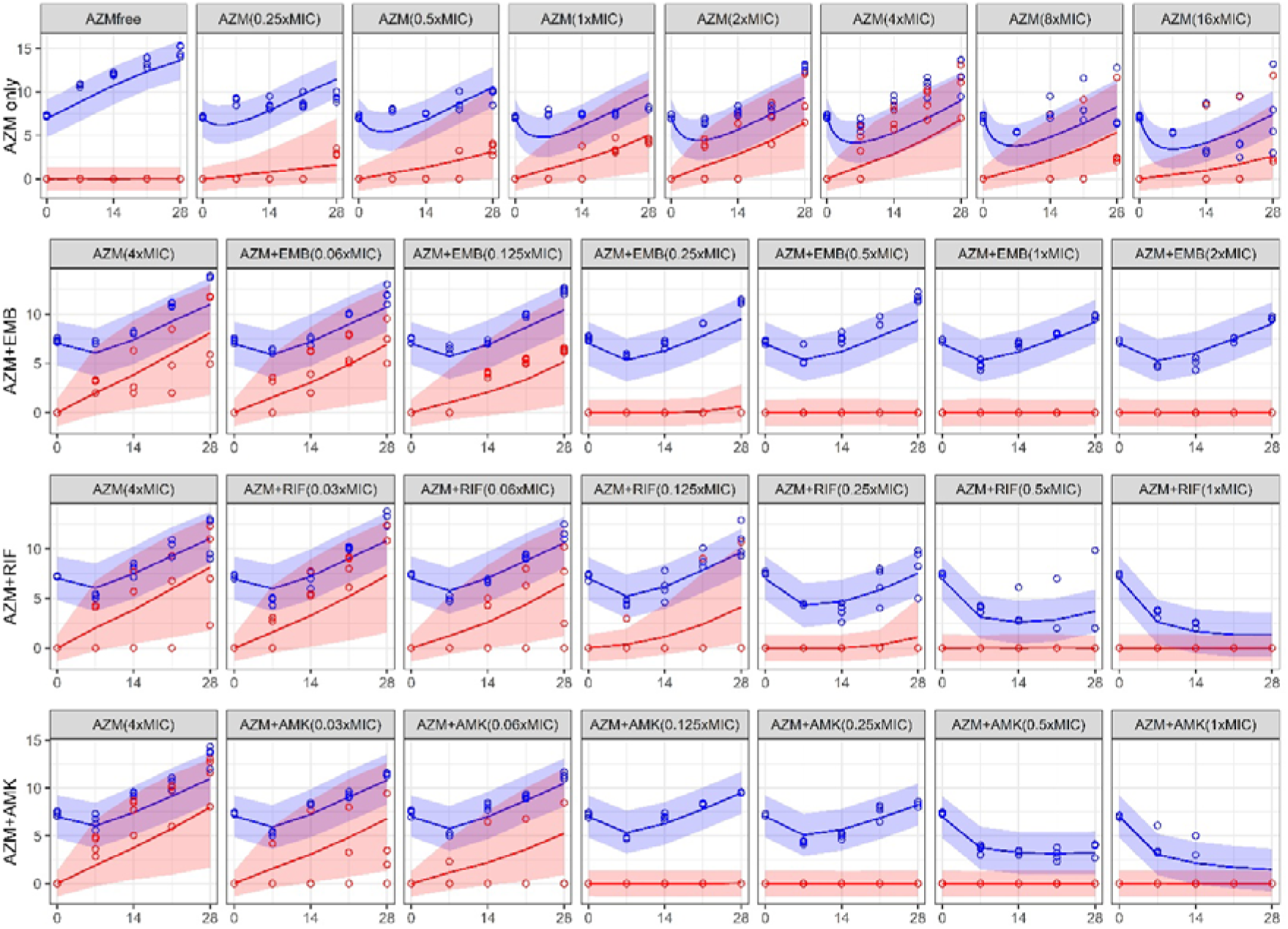
Observed and pharmacodynamic (PD)-model predicted bacterial counts. Y-axis represents the bacterial count (logLL CFU/mL), and the X-axis represents the time in days. “AZM only” represents the time-kill assay (TKA) results with azithromycin (AZM) monotherapy, whereas “AZM+EMB,” “AZM+RIF,” and “AZM+AMK” represent the results of combination TKAs in which each antimicrobial was co-incubated with AZM at 4× MIC over a concentration range of 0.03× to 2× MIC. Total bacteria are presented in blue, and resistant bacteria are presented in red. Observed values are represented by circles, model-predicted means by solid lines, and the 90th percentile of the model predictions by shaded areas.

### PK/PD Model Simulation

The simulated resistance and bactericidal rates of AM under various dosing conditions for the two-drug combinations AZM+EMB, AZM+RIF, and AZM+AMK are shown in Figures 2, 3, and 4. For all concomitant agents, daily dosing regimens were less likely to induce resistance than a three-times weekly regimen. Among the daily regimens, the resistance rates were lower with twice-daily dosing than those with once-daily dosing, even for the same total daily dose. EMB was the most potent in suppressing the emergence of resistance, although it was simulated to have no bactericidal effects (Figure 2). Furthermore, in the three-times-weekly dosing regimen, RIF was simulated to not affect resistance suppression (Figure 4). Figure 5 shows the simulated additive effects of RIF, AMK, or RIF+AMK on the AZM+EMB regimen using the guideline-recommended doses as the baseline. The combined effects of AMK and RIF on the suppression of macrolide resistance were minimal. In contrast, for bactericidal activity, both RIF and AMK were simulated to provide additional effects on EMB.

**Figure 2.**
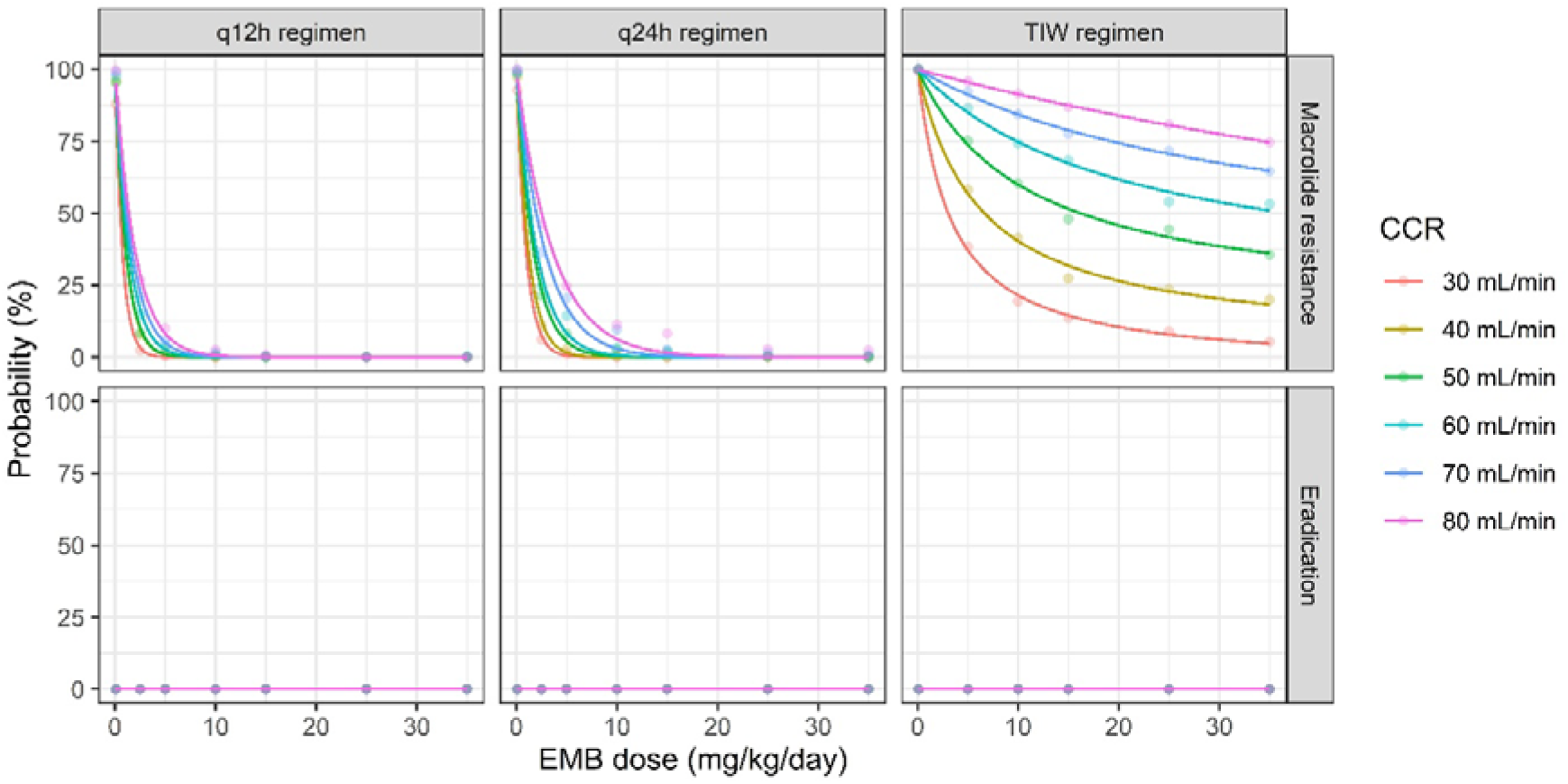
Simulated macrolide resistance and bactericidal rates by ethambutol (EMB) dosages. Graphs are arranged by dosing regimen (twice daily [q12h], once daily [q24h], and thrice weekly [TIW]) in columns, with colours indicating different levels of creatinine clearance (mL/min). Dots represent simulated values for each dose, and lines represent smoothed curves. AZM dose was fixed at 250 mg/day for daily administration and 500 mg/day for the TIW regimen.

**Figure 3.**
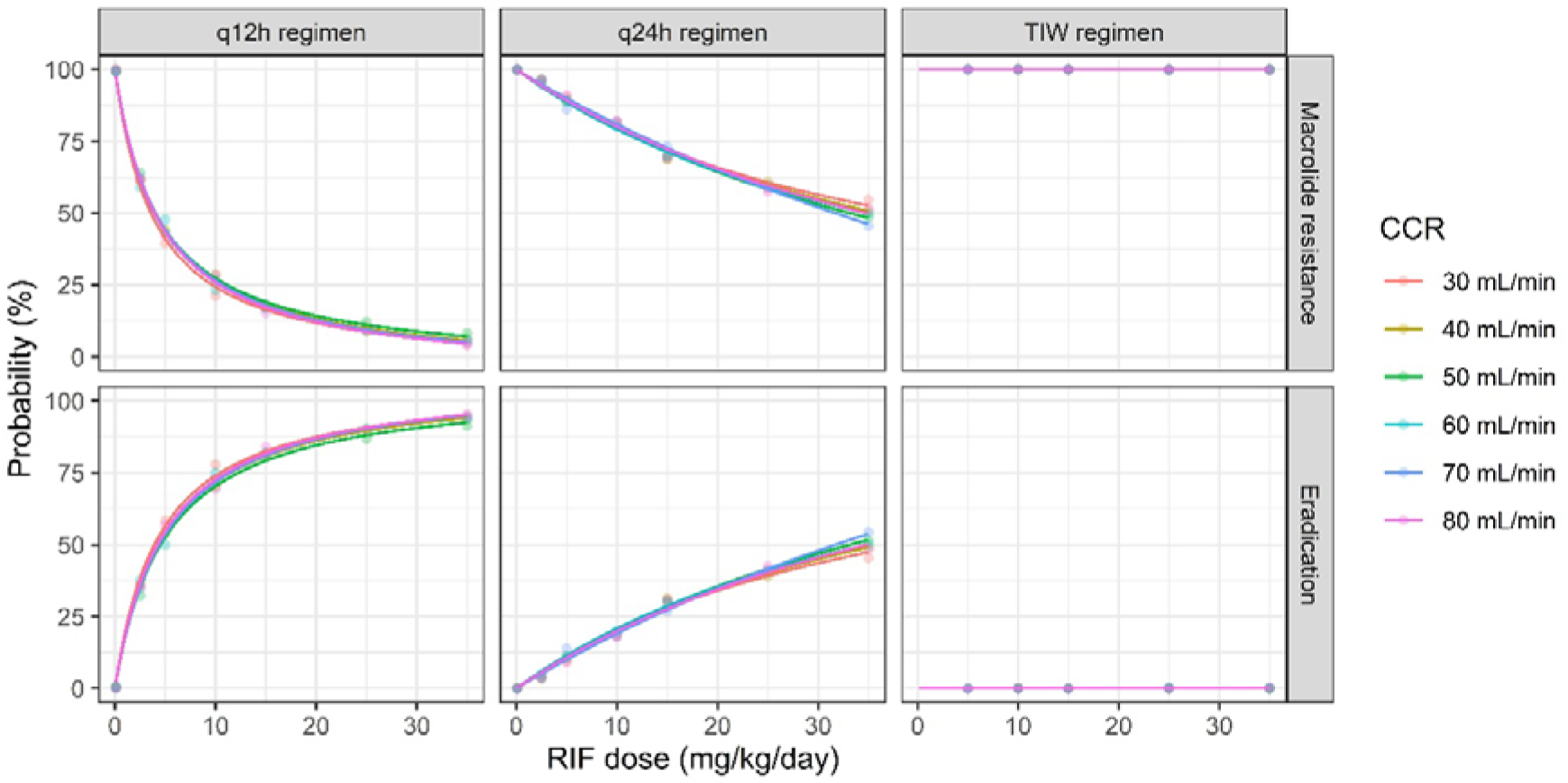
Simulated macrolide resistance and bactericidal rates by rifampicin (RIF) dosages. Simulation conditions were identical to those described in Figure 2, except that RIF was used instead of EMB.

**Figure 4.**
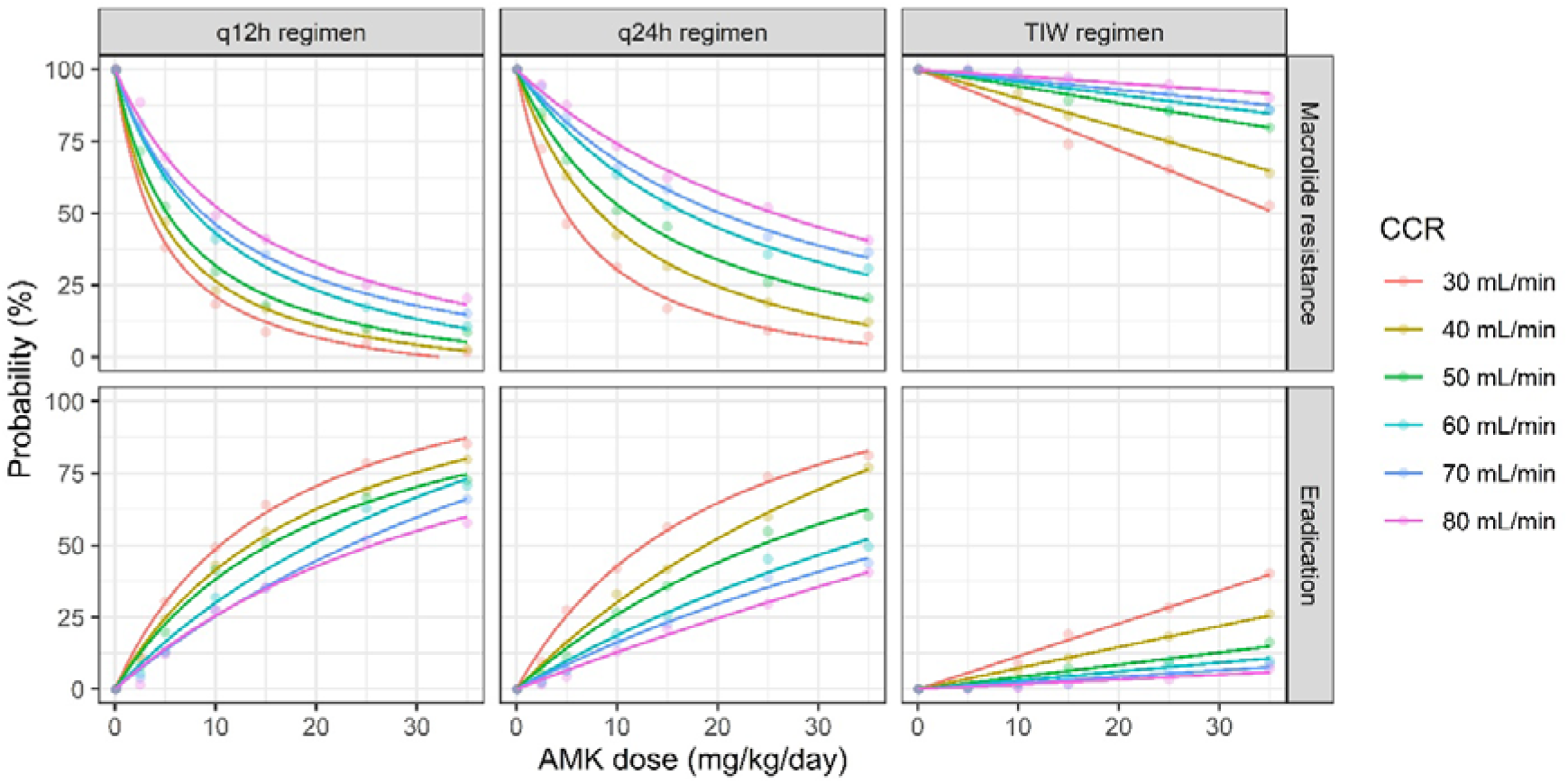
Simulated macrolide resistance and bactericidal rates by amikacin (AMK) dosages. Simulation conditions were identical to those described in Figure 2 except that AMK was used instead of EMB.

**Figure 5.**
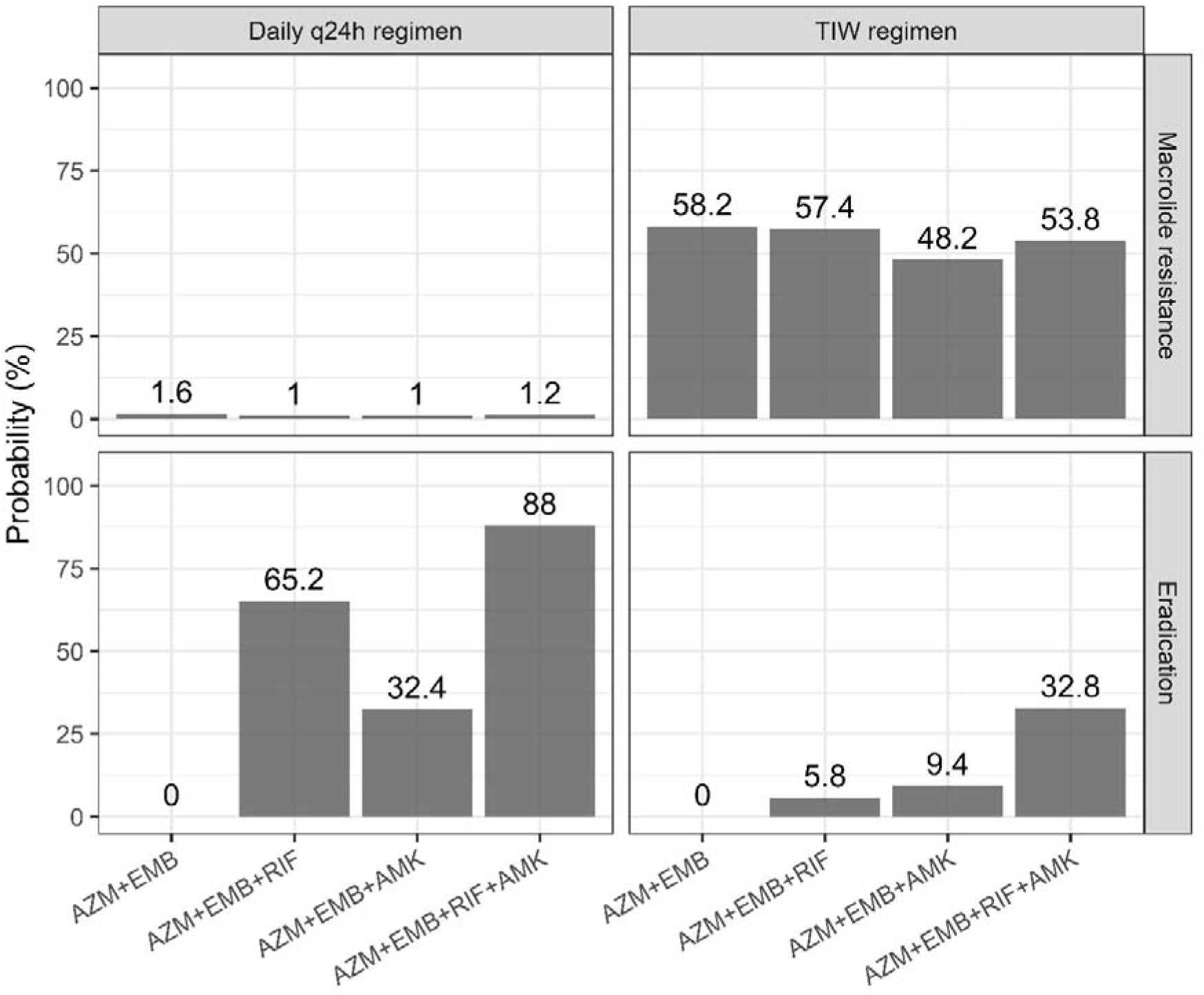
Simulated macrolide resistance and bactericidal rates under guideline dosing regimens. TIW, three times weekly; Simulations were performed under the following conditions: AZM 250 mg once daily or 500 mg TIW; EMB 15 mg/kg once daily or 25 mg/kg TIW; RIF 10 mg/kg once daily or 600 mg TIW; and AMK 15 mg/kg once daily or 25 mg/kg TIW.

## DISCUSSION

To the best of our knowledge, this is the first study to identify the concentrations of concomitant antimicrobial agents required to suppress macrolide resistance in *M. avium*. This is also the first study to quantitatively evaluate, through PK/PD analysis, the relationship between antimicrobial dosing regimens and resistance emergence. Furthermore, this study proposes a novel TKA methodology designed to assess the resistance-suppressive effects of combination therapy, representing a methodological advancement in the *in vitro* studies of multidrug regimens.

In this study, macrolide resistance most frequently emerged at 2–4× the MIC of AZM, whereas it was less frequent at 8–16× the MIC (Figure 1 and S1). This trend was also evident in our preliminary experiments, wherein CLR was used instead of AZM (data not shown). Since macrolides exhibit time-dependent bactericidal activity,^33^ their killing effect per unit time is considered constant regardless of the concentration. Consistent with this characteristic, no marked reduction in bacterial counts was observed at 8–16× MIC; however, the relatively low frequency of resistance emergence at these concentrations may be attributed to approaching the mutant prevention concentration.^34^ In addition to the high-concentration range, resistance emergence was less likely at concentrations below the MIC. *M. avium* is known to upregulate efflux pumps following AZM exposure,^31^ which may reduce intracellular macrolide concentrations and consequently decrease the possibility of resistance development. The finding that the concomitant agents suppressed macrolide resistance, even at sub-MIC concentrations, was particularly interesting (Figure 1 and S2–4).

The PK/PD model developed in this study integrates AM concentrations, enabling the simulation of resistance emergence, which reflects actual drug exposure at pulmonary infection sites. This approach has direct implications for the design of clinically relevant dosing regimens. Consistent with previous reports, EMB exhibited the strongest resistance-suppressive effect, whereas RIF showed minimal effect.^8,35^ Since EMB did not exhibit *in vitro* bactericidal activity, it did not contribute to bacterial reduction in the PK/PD simulations. However, since the present analysis did not incorporate host immune responses into the model, these results do not imply that the AZM + EMB regimen failed to achieve sputum culture conversion in clinical settings. Moreover, twice-daily dosing resulted in lower resistance rates than those with once-daily dosing, even at the same total daily dose. Considering that EMB, RIF, and AMK are associated with dose-dependent adverse effects, such as optic neuropathy,^36^ hepatotoxicity,^37^ gastrointestinal symptoms,^5^ and ototoxicity,^38^ dividing the total daily dose into twice-daily administration is a potential strategy to reduce toxicity while maintaining resistance suppression.

Although CFZ showed a rapid decline in the MB7H9b concentration (Figure S4), it presumably resulted from adsorption to the plastic rather than chemical degradation. Polypropylene bottles were used in this study; a supplementary experiment using polystyrene 96-well plates demonstrated that CFZ concentrations remained stable over 28 d at 37°C. Considering these complex physicochemical properties, the actual CFZ concentration to which bacteria are exposed cannot be reliably determined. Furthermore, as the CFZ concentrations in the ELF and AM have not been established, it is not feasible to include CFZ in PD modelling or PK/PD simulations.

This study had some limitations. We used an *in vitro* model, which did not account for host factors such as immune responses. Furthermore, the PK model, particularly the parameters describing drug penetration into the pulmonary ELF and AM, were based on data derived from a limited number of participants, which may have restricted the external validity of the simulations. Furthermore, the present simulations of three- and four-drug combinations were based on data obtained from two-drug experiments, potentially underestimating the synergistic effects that could occur with regimens containing three or more agents. Finally, this experiment was conducted using a single ATCC strain. Therefore, validation studies using clinical isolates are required in the future. Despite these limitations, this study provides a novel framework for quantitatively evaluating the relationship between antimicrobial dosing regimens and suppression of macrolide resistance in MAC pulmonary disease, offering valuable insights for optimizing clinical combination therapy.

## Acknowledgments

Part of this work was presented at the 8^th^ World Bronchiectasis Conference, held in Brisbane, Australia on 14–17 July 2025. We thank Kodai Ikeda, Nanaka Arai, Tsumugi Umezawa, and Haruto Ozawa (Meiji Pharmaceutical University) for their technical assistance with TKA experiments.

We would like to thank Editage (www.editage.jp) for English language editing.

## Author Contributions

Fumiya Watanabe: Conceptualization, Methodology, Investigation, Data curation, formal analysis, Software, Validation, Visualization, Writing of the original draft. Yuta Morishige: Methodology, Investigation, Resources, Writing – Review and Editing. Akio Aono: Conceptualization, Methodology, Resources, Writing – review, and editing. Yuji Morita: Methodology, Resources, Project Administration, Writing – Review, and Editing. Kozo Morimoto: Project administration, Supervision, Writing, reviewing, and editing. Satoshi Mitarai: Methodology, Project administration, Resources, Supervision, Writing, review, and editing. Kazuhiko Hanada: Conceptualization, Methodology, Supervision, Software, Writing – review, and editing. All authors have approved the final version of the manuscript for publication.

## Declaration of generative AI and AI-assisted technologies in the manuscript preparation process

No generative AI or AI-assisted technologies were used in preparing this manuscript.

## Ethics Statement

This study did not involve human or animal participants; therefore, ethical approval was not required.

## Data Availability Statement

The data that support the findings of this study are available from the corresponding author upon reasonable request.

