## Supplementary materials for "Concentration-dependent suppression of macrolide resistance in *Mycobacterium avium* by combination therapy: an improved *in vitro* time-kill assay and PK/PD modelling study"

Text S1. Pharmacodynamic model

The pharmacodynamic (PD) model was developed using NONMEM version 7.6.0 (ICON Development Solutions, Hanover, MD, USA) and the ADVAN13 subroutine. Values below the lower limit of detection (100 CFU/mL) were fitted using Beal's M3 method.[1] A two-compartment model, including a macrolide-susceptible and -resistant bacterial population, was fitted to the log-transformed (i.e., $\log_{10}$ CFU/mL) time-kill assay data. In the initial conditions, the susceptible compartment was set to the inoculum level (approximately 7 $\log_{10}$ CFU/mL), whereas the resistant compartment was set to zero. A schematic illustration of this process is provided below.

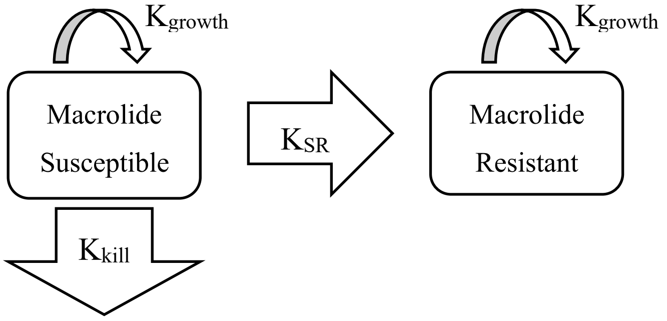

where $K_{growth}$, $K_{kill}$, and $K_{SR}$ are the bacterial growth rate constant, bacterial killing rate constant, and macrolide resistance acquisition rate constant, respectively. The $K_{growth}$ is represented by Eq. 1.

Eq. 1. $K_{growth} = K_g \times (1 - \frac{Susceptible + Resistant}{N_{max}})$

where $N_{max}$ is the maximum bacterial capacity of the system (16 $\log_{10}$ CFU/mL). The effects of azithromycin (AZM) and combination drugs such as ethambutol (EMB), rifampicin (RIF), and amikacin (AMK) on bacterial reduction were determined using the sigmoid-maximum effect (sigmoid-Emax) model in Eq. 2.

Eq. 2. $K_{kill} = \frac{K_{AZM,max} \times AR \times Conc_{AZM}{}^{hill1}}{K_{AZM,50}{}^{hill1} + Conc_{AZM}{}^{hill1}} + \frac{K_{Comb,max} \times Conc_{Comb}{}^{hill\ 3\ to\ 5}}{K_{Comb,50}{}^{hill\ 3\ to\ 5} + Conc_{Comb}{}^{hill\ 3\ to\ 5}}$

where $K_{max}$, $K_{50}$, Conc, and Comb are the maximum bacterial killing rate constant, concentration required to produce half of the maximum killing rate, concentration of each drug, and combination of drugs, respectively. In this study, the effect of the combination drugs on bacterial reduction was expressed in an additive manner since only one macrolide-plus-single-agent condition was experimentally evaluated; subsequent simulations involved scenarios with three to four concomitant agents. An additive model was used to avoid overestimating bacterial killing owing to unforeseen synergistic effects in the simulated multidrug combinations. AR refers to adaptive resistance to macrolides observed after day 7 of the time-kill assay, which occurs independently of mutations in the *rrl* gene. Since AZM exposure led to a 56-fold increase in the expressions of *MAV_3306* and *MAV_1406*, which encode an ABC transporter and a major facilitator superfamily efflux pump in *Mycobaterium avium*, respectively, low-level adaptive resistance (AR) was observed,[2] which is consistent with previous reports and with the findings of the present study. The AR was modelled using Eq. 3.

Eq. 3. $\mathrm{AR} = 1 - AR_{max} \times (1 - e^{-K_{delay} \times Time})$

where $AR_{max}$ and $K_{delay}$ represent the maximum adaptive resistance effect and the lag time required to reach that effect, respectively. The $K_{SR}$ was modelled using the asymmetric sigmoid-$E_{max}$ model, as shown in Eq. 4.

Eq. 4. $K_{SR} = \frac{K_{SR,max} \times {Conc_{AZM}}^{hill2}}{{K_{SR,50}}^{hill2} + {Conc_{AZM}}^{hill2}} \times e^{-\lambda \times Conc_{AZM}} \times (1 - \alpha_{comb} \times Conc_{Comb})$

where $K_{SR,max}$, $K_{SR,50}$, λ, and α were maximum macrolide resistance acquisition rate constant, concentration needed to produce one-half of the maximum resistance rate, asymmetry coefficient, and coefficient describing the concentration-dependent suppression coefficient of combination drugs, respectively. To prevent negative values, the term $(1 - \alpha_{comb} \times Conc_{Comb})$ was bounded at zero.

The concentration parameters of each antimicrobial incorporated into the PD model were normalized to their respective MIC values and expressed as multiples of the MIC.

The stability of the considered antimicrobial agents in Middlebrook 7H9 broth at 37°C was evaluated. Quality control samples at 0.25× and 16× MIC for each drug were prepared in quadruplicate and incubated with aliquots collected at 0, 7, 14, 21, and 28 d after initiation. Drug concentrations were measured using validated high-performance liquid chromatography-mass spectrometry, as described previously.[3] The first-order degradation rate constants ($K_{deg}$) were calculated for each drug. The antibiotic concentrations used in the PD model during the time-kill assay were corrected based on these $K_{deg}$ values.

For the random effect of the PD model, the inter-batch variability of $K_{SR}$ was incorporated using an exponential error model, and the residual variability for the resistant subpopulation and total bacteria was described using an additive error model.

The reliability of the final PD model was assessed using goodness-of-fit plots, visual predictive checks, and bootstrap analyses.

Text S2. Pharmacokinetic profiles of the antimicrobial agents in the epithelial lining fluid and alveolar macrophages

To simulate the risk of macrolide resistance under different dosing regimens, the pharmacokinetic (PK) profiles of each drug in the epithelial lining fluid (ELF) and alveolar macrophages (AM) were incorporated into a PK model based on previously published data. Since protein binding in ELF is minimal, drug concentrations measured in ELF are assumed to reflect entirely unbound drugs.[4] The post-antibiotic effects (PAE) of EMB, RIF, and AMK against *M. avium* in the presence of macrolides have been reported as 13.8 ± 1.5 h, 24.0 ± 3.9 h, and 10–20 h, respectively.[5,6] However, these PAE were observed upon the first exposure to macrolides and may not reflect the conditions during prolonged chemotherapy, as in MAC-PD. *M. avium* upregulates efflux pumps 3–5 d after AZM exposure,[2] and efflux pump overexpression is known to reduce PAE.[7] Therefore, as the PAE of *M. avium* following macrolide exposure under prolonged treatment conditions remains unclear, it was not included in the model.

AZM conditions

PK data for AZM in the ELF and AM are available from three studies,[8–10] all of which used a loading dose of 500 mg, followed by 250 mg once daily for 4 d. In each study, ELF concentrations ranged from approximately 1–3 mg/L, whereas AM concentrations were substantially higher at approximately 50–300 mg/L, and these concentrations were sustained for at least 124 h post-dose. Accordingly, for both the 250-mg once-daily and 500-mg three-times-weekly regimens, the AZM concentrations in the ELF and AM were assumed to remain within these respective ranges. The pH within AM is acidic,[11] which reduces macrolide activity to approximately 30–50% of that observed under neutral conditions.[12] Accordingly, AZM activity in the AM compartment was modelled by applying a one-third correction factor to the AM concentration. Coadministration with RIF decreases the steady-state AUC of AZM by approximately 24%[13]; therefore, a correction factor of 0.76 was applied to AZM concentrations in simulations involving RIF. The MIC distribution was modelled as log-normal, with AZM MICs assumed to be centred at 8, 16, and 32 mg/L, in accordance with reported values.[14]

EMB conditions

The MIC distribution of EMB was modelled as log-normal according to previously reported values.[15] PK data describing EMB concentrations in the plasma, ELF, and AM were derived from a cohort of 157 patients with tuberculosis in Malawi.[16] The median age was 37 years (interquartile range [IQR], 28–39 years), 76.4% were male, and the median body mass index was 18.4 kg/m² (IQR, 17.0–19.8). The administered EMB doses were 550 mg (1.9%), 825 mg (68.2%), 1100 mg (28.7%), and 1375 mg (1.3%). Plasma samples were collected prior to dosing and at 0.5, 1, 2, 3, 4, 5, 6, and 8 h. ELF and AM samples were obtained at 2, 4, and 6 h post-dosing. The final population PK parameter estimates are presented in Table S1.

Table S1. PK parameter estimates of EMB

| | Mean | Inter-individual variability<br>Eta value (CV) |
|---|---|---|
| CL/F [L/h] | 43.0 | 0.098 (29.8%) |
| V/F [L] | 401.0 | 0.063 (25.1%) |
| Ka [/h] | 0.36 | 0.463 (68.0%) |
| CCR on CL/F | 0.12 | |
| $R_{ELF/total\ plasma}$ | 4.0 | 0.139 (48.3%) |
| $R_{AM/total\ plasma}$ | 15.0 | 0.204 (43.3%) |

CL/F, apparent clearance; V/F, apparent volume of distribution; Ka, absorption rate constant; CCR, creatinine clearance; $R_{ELF\ /\ total\ plasma}$; ELF/plasma ratio; $R_{AM/\ total\ Plasma}$, AM/Plasma ratio; CV, coefficient of variation.

RIF conditions

The MIC distribution of RIF was modelled as log-normal in accordance with reported values.[15] PK data describing RIF concentrations in plasma, ELF, and AM were derived from the same cohort as EMB.[16] The final population PK parameter estimates of RIF are shown in Table S2.

Table S2. PK parameters of RIF

| | Mean | Inter-individual variability<br>Eta value (CV) |
|---|---|---|
| CL/F [L/h] | 12.2 | 0.028 (16.7%) |
| V/F [L] | 22.2 | 0.382 (61.8%) |
| Ka [/h] | 0.24 | 0 fixed |
| $R_{ELF/total\ plasma}$ | 1.97 | 0.219 (46.8%) |
| $R_{AM/total\ plasma}$ | 1.35 | 1.03 (101.5%) |

CL/F, apparent clearance; V/F, apparent volume of distribution; Ka, absorption rate constant; $R_{ELF\ /\ total\ plasma}$, ELF/plasma ratio; $R_{AM/\ total\ Plasma}$, AM/Plasma ratio; CV, coefficient of variation

AMK conditions

The MIC distribution of AMK was modelled as log-normal according to previously reported values.[17] The PK parameters of AMK were obtained from a cohort of 137 older patients from South America.[18] The median age of the included patients was 76 years (range, 65–93), the mean ± standard deviation creatinine clearance was 86.6 ± 40.3 mL/min, and the median AMK dose was 14 mg/kg/day (range, 5.7–22.6). Serum AMK concentrations were sampled at an average of three time points per patient, between 0.5 and 77 h post-after administration. The parameter estimates for the final population PK model are presented in Table S3.

Table S3. PK parameters of AMK

| | Mean | Inter-individual variability Eta value (CV) |
|---|---|---|
| CL [L/h] | 12.2 | 0.028 (16.7%) |
| Vc [L] | 22.2 | 0.382 (61.8%) |
| Q [L/h] | 0.5 | 0 fixed |
| Vp [/h] | 11.9 | 0 fixed |
| CL * CCR/80 $^{THETA}$ | 0.7 | |
| Vc * BMI/25 $^{THETA}$ | 1.16 | |
| $R_{ELF/total\ plasma}$ | 0.50 | 0.84 (91.7%) |
| $R_{AM/total\ plasma}$ | N.A. | N.A. |

CL, clearance; Vc, central volume of distribution; Q, intercompartment clearance; Vp, peripheral volume of distribution; CCR, creatinine clearance; BMI, body mass index; $R_{ELF\ /\ total\ plasma}$, ELF/plasma ratio, $R_{AM/\ total\ Plasma}$, AM/Plasma ratio; N.A., not available; CV, coefficient of variation.

Although the penetration ratio of AMK into the ELF is approximately 0.5 of the plasma concentrations, with 91.7% inter-individual variability (CV),[19] to our knowledge, no data are available regarding its penetration into the AM. Therefore, AM penetration was assumed to be equivalent to that in the ELF.

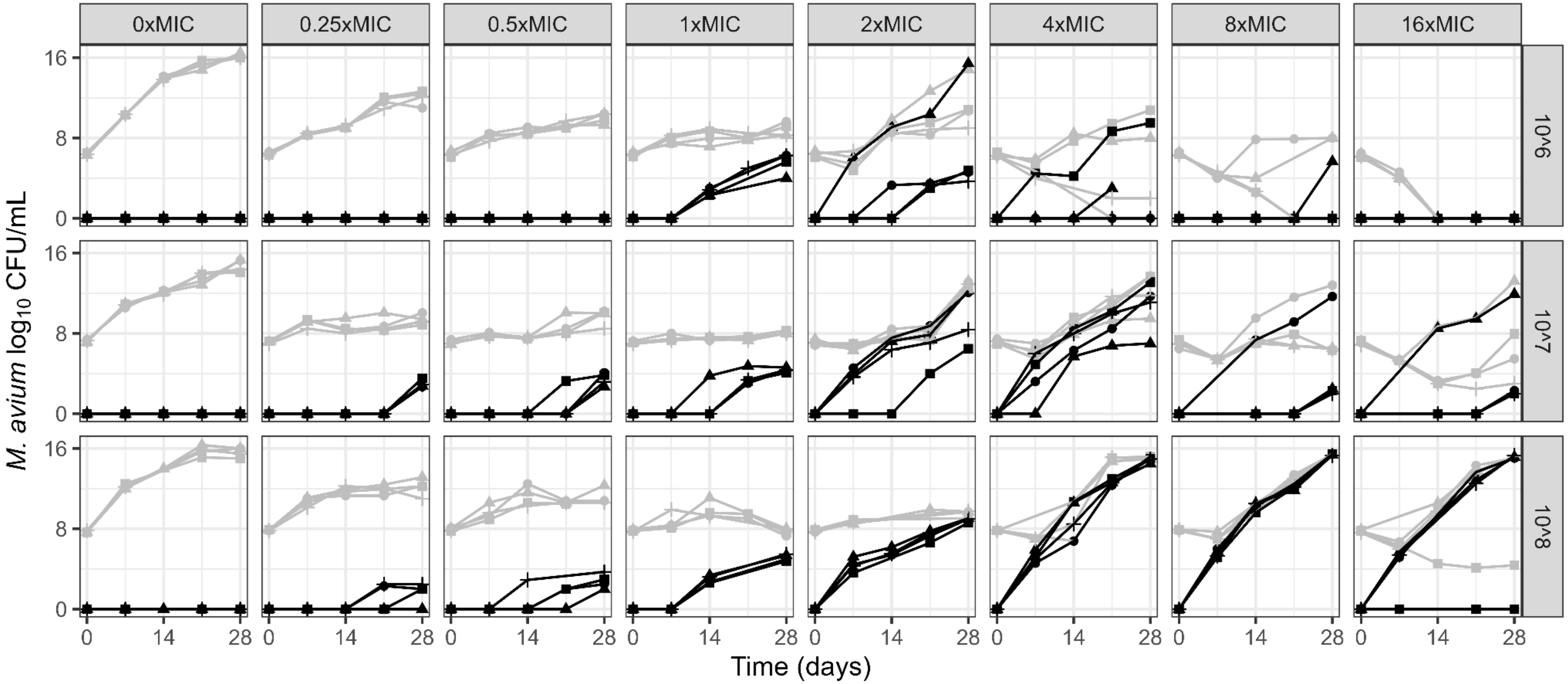

Figure S1. TKA results for macrolide monotherapy

Y-axis represents the logarithmic *M. avium* concentration and X-axis indicates the time elapsed after the start of incubation. Each experiment was performed in quadruplicate. Grey dots and lines represent the total bacterial counts, whereas black dots and lines represent the counts of resistant bacteria. Column labelled "0×MIC" indicates the growth control condition without AZM, and results are shown for cultures with AZM concentrations ranging from 0.25× to 16× MIC. The numbers $10^6$, $10^7$, and $10^8$ in each row denote the initial inoculum levels, corresponding to $2\times10^6$, $2\times10^7$, and $2\times10^8$ CFU/mL, respectively.

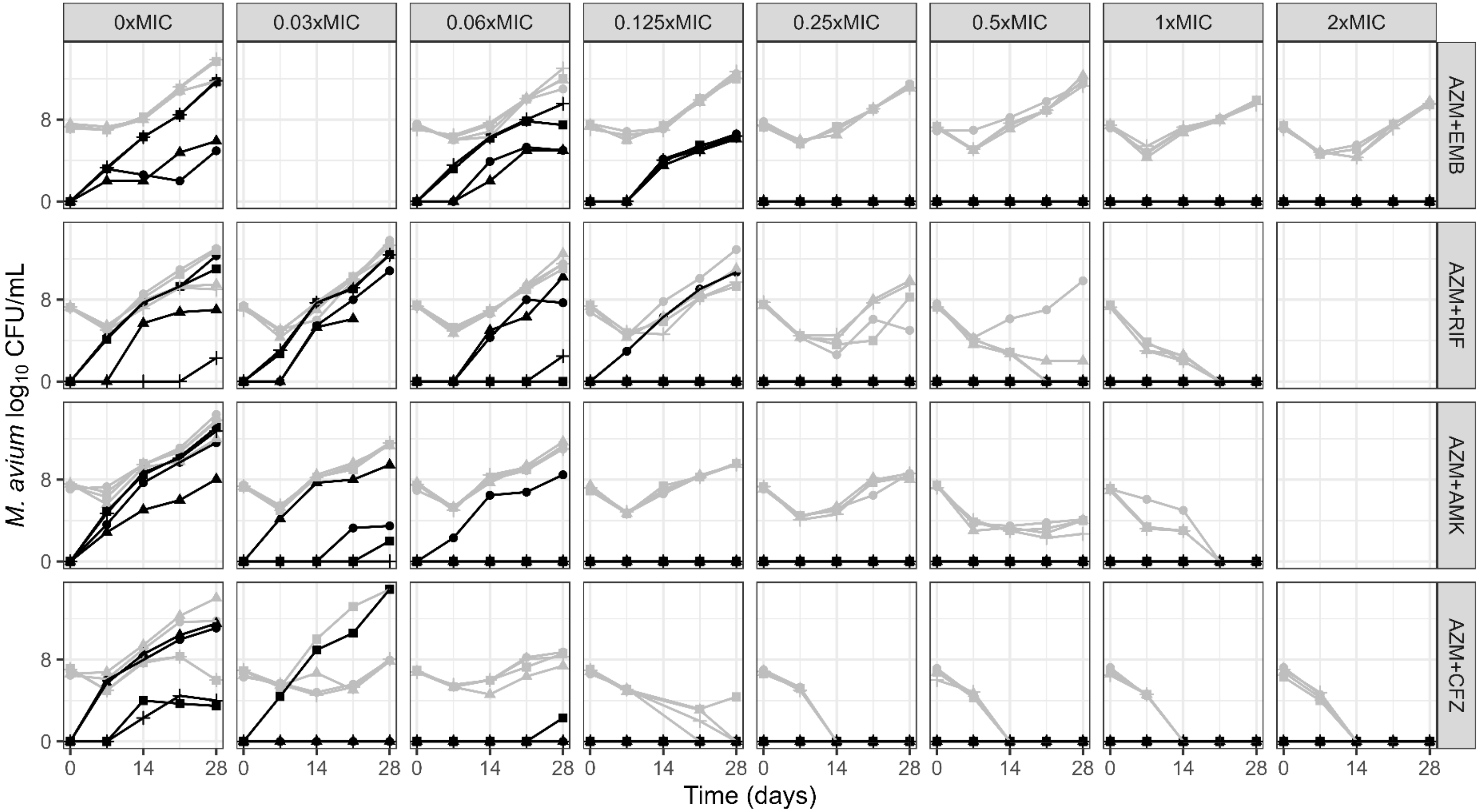

Figure S2. TKA results under combination exposure (initial inoculum: $10^7$ CFU/mL).

EMB, ethambutol; RIF, rifampicin; AMK, amikacin; CFZ, clofazimine. Grey indicates the time course of total bacterial counts, whereas black indicates the time course of resistant bacterial counts. All conditions included AZM at 4× MIC and the column concentrations represented those of the concomitant agents. Initial bacterial burden was $2×10^7$ CFU/mL.

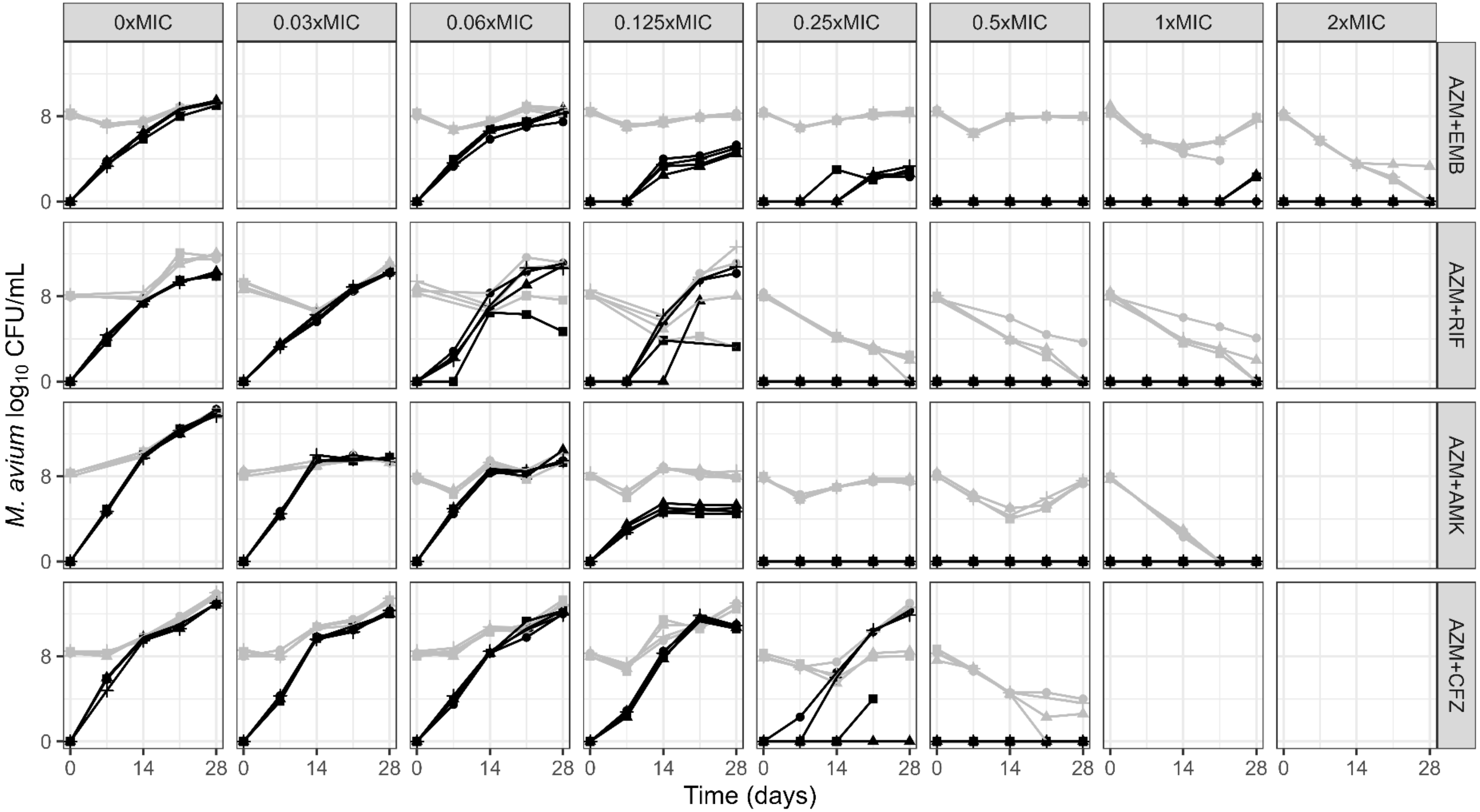

Figure S3. TKA results under combination exposure (initial inoculum: $10^8$ CFU/mL).

EMB, ethambutol; RIF, rifampicin; AMK, amikacin; CFZ, clofazimine. Grey indicates the time course of total bacterial counts, whereas black indicates the time course of resistant bacterial counts. All conditions included AZM at 4× MIC and the column concentrations represented those of the concomitant agents. Initial bacterial burden was $2\times10^8$ CFU/mL.

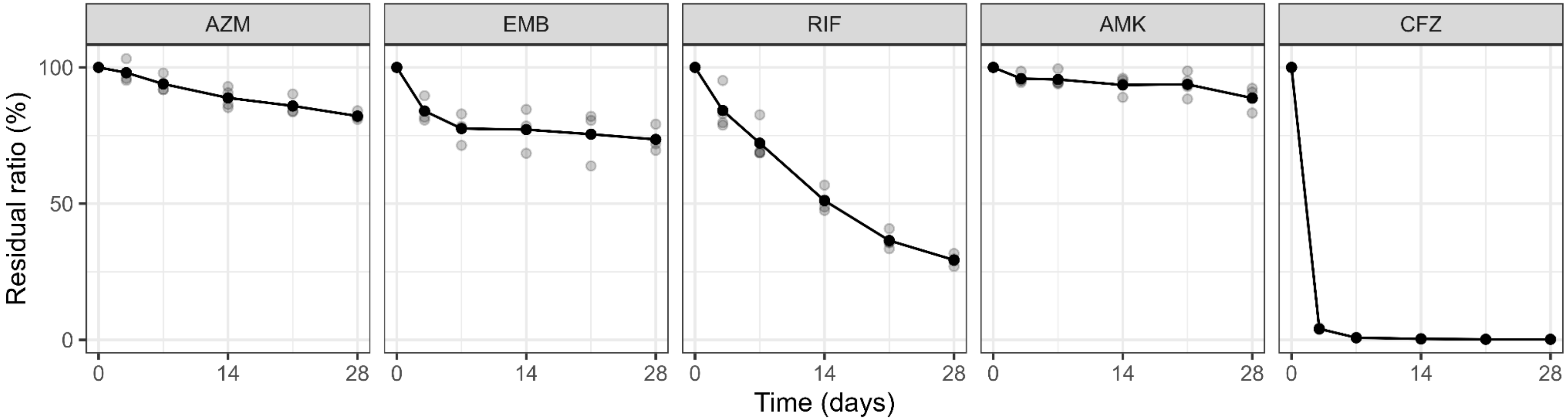

Figure S4. Residual Ratios of Antimicrobial Agents in the Culture Medium

Drugs stability in MB7H9b with 0.05% Tween 80 and 10% OADC without bacteria at 37℃ until 28 days (N = 4). Grey dots represent the observed values, and black dots and lines represent the mean values.

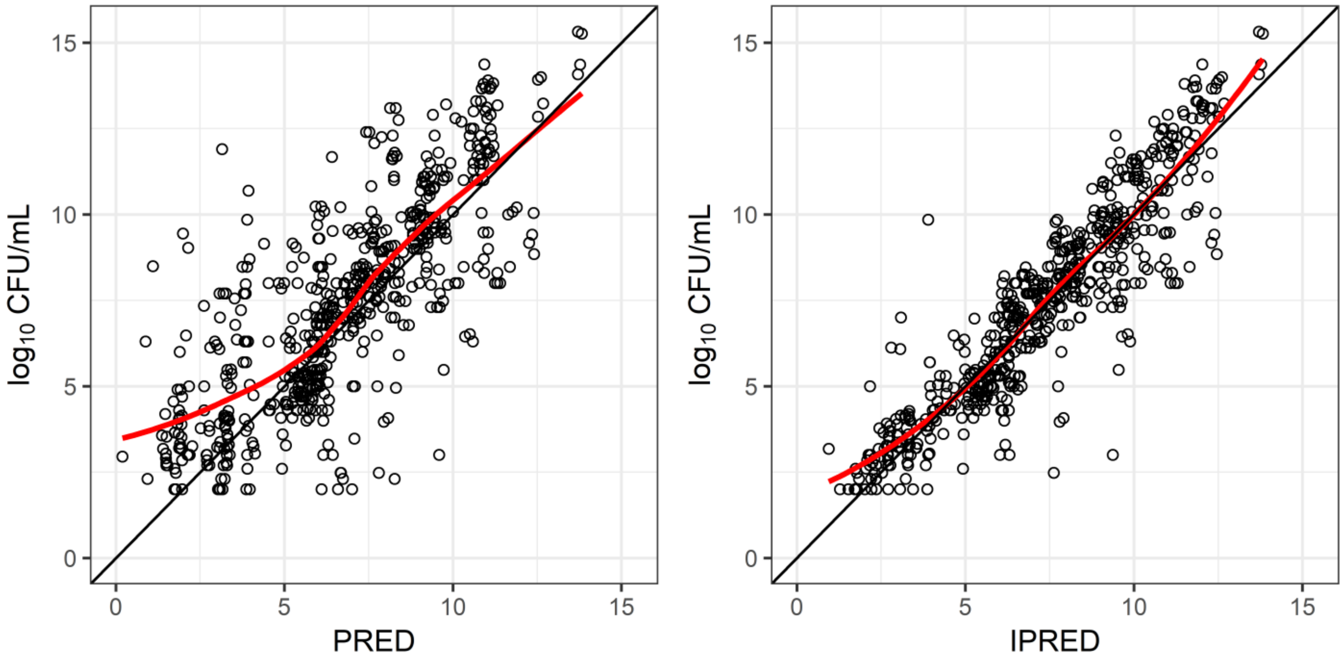

Figure S5. Goodness-of-fit plot of the PD model.

Y-axis represents the observed $log_{10}$ values, and the X-axis represents the model-predicted values. PRED indicates population-predicted mean values, and IPRED indicates individual-predicted values.
